# Protein Kinase A Activity In The Leading Edge of Migrating Cells Is Dependent On The Activity of Focal Adhesion Kinase

**DOI:** 10.1101/2022.09.17.508387

**Authors:** Kathryn V. Svec, Mingu Kang, Alan K. Howe

**Affiliations:** Department of Pharmacology, University of Vermont, Burlington, VT, United States; Department of Molecular Physiology and Biophysics, University of Vermont, Burlington, VT, United States; University of Vermont Cancer Center, University of Vermont, Burlington, VT, United States

## Abstract

Protein Kinase A (PKA) is a pleiotropic serine/threonine kinase whose localized and dynamic activity is required for cellular migration. Several cell types have shown robust PKA activity in the leading edge during migration. This activity is regulated by changes in actomyosin contractility, but the mechanism of the mechanochemical regulation of PKA remains unclear. Our ongoing investigation into this mechanism led us to discover a novel relationship between PKA and the non-receptor tyrosine kinase Focal Adhesion Kinase (FAK). Here, we find that while inhibition of actomyosin contractility leads to a relatively slow decrease in total cellular levels of phosphorylated FAK over tens of minutes, it decreases the focal adhesion-associated pool of both phospho-FAK and phospho-paxillin within two minutes, similar to the timing of the effect of inhibition of contractility on leading edge PKA activity. We then show that pharmacologic inhibition of FAK rapidly decreases leading edge PKA activity. Additionally, leading edge PKA activity is significantly reduced in FAK-null mouse embryonic fibroblasts and essentially eliminated by also silencing expression of the closely-related kinase Pyk2 in these cells. Importantly, leading edge PKA activity in FAK-null cells is rescued by re-expression of wild-type but not kinase-dead FAK. Taken together, these observations show that FAK activity is required for localized regulation of PKA during cell migration and, in so doing, establish a novel and unexpected connection between an adhesion-regulated tyrosine kinase and a serine/threonine kinase canonically downstream of G-protein coupled receptor and second-messenger signaling.

## INTRODUCTION

Cell migration is essential to many diverse normal and pathological processes alike, including wound healing, development, and cancer cell metastasis. Many decades of research have elucidated myriad signaling pathways involved in cell migration, but in such an intricate process, seemingly discrete signaling nodes must work in concert to coordinate directed motility.

Substantial evidence points to Protein Kinase A (PKA) as an important mediator of cell migration upstream of membrane dynamics, actin organization, and many other processes critical to cell motility [1-3]. PKA is a principle serine/threonine kinase canonically activated downstream of G-protein coupled receptor (GPCR) activation and the activation of adenylyl cyclases [4]. Further, we and others have shown that PKA is active in the leading edge of migrating cells [5-11], alongside innumerable other signaling molecules that guide and coordinate cell migration, such as Cdc42 and RhoA [12, 13]. Our previous work identified a dependence of leading edge PKA activity on cellular contractility as it is rapidly lost upon treatment with myosin II inhibitor, blebbistatin [11]. Further, durotactic stretch can potentiate rapid, local bursts of PKA activity and PKA is required for mechanically gated cell migration, or durotaxis [11]. Thus, PKA activity in the leading edge of migrating cells is coupled to cellular contractility.

However, it remains unclear how PKA activity is regulated by mechanical inputs. Generally, cells sense changes in extracellular forces via focal adhesion complexes and associated dynamic actin structures. Mechanical force is translated into chemical signals via physical deformation of focal adhesion components, allowing for the confluence of different combinations of proteins and modified signaling [14-17]. Prominent among mechanically sensitive signaling proteins is Focal Adhesion Kinase (FAK), a nonreceptor tyrosine kinase well-characterized for its role in adhesive signaling, principally through the phosphorylation of paxillin and several other focal adhesion proteins [18, 19]. Further, FAK is involved in the iterative probing of the extracellular environment performed by focal adhesions through fluctuations in force during durotaxis [20] and is a key mediator of signaling to and from contractile forces [21-24]. Yet while FAK mediates mechanochemical signaling and is required for durotaxis[20, 25], it has never been implicated in the regulation of PKA activity, in any context.

Several studies provide evidence of crosstalk between tyrosine kinases and PKA signaling, but almost exclusively involving the regulation of tyrosine phosphorylation by PKA [26-34]. Though it is uncommon, it is not unheard of for a tyrosine kinase to regulate PKA activity. Both receptor tyrosine kinases and Src family kinases have been shown to regulate PKA activity by phosphorylating the PKA catalytic subunit directly [35, 36]. However, this work has been largely conducted *in vitro* and has not yet extended to FAK. Therefore, given the regulation of leading edge PKA activity by mechanical inputs and the central role of FAK as a mechanotransducer, we tested whether FAK plays a role in regulating localized PKA activity in migrating cells.

## RESULTS

### Disrupting actin polymerization does not lead to an immediate loss of leading edge PKA activity

Our prior research identified leading edge PKA activity as being rapidly modulated by mechanical inputs, both applied and intrinsic. Transduction of mechanically-gated signals during migration depends on actin polymerization [37, 38]. Therefore, we examined PKA activity in migrating cells before and after treatment with inhibitors of actin polymerization. Specifically, PKA activity was monitored in migrating SKOV-3 ovarian cancer cells expressing a membrane targeted version of a FRET-based biosensor sensitive to rapid and dynamic changes in PKA activity (pm-AKAR3) [39]. Upon phosphorylation by PKA, the single-chain biosensor undergoes a reversible conformational change, causing increased energy transfer between the ECFP and cpVenus [39]. By calculating the ratio of these two fluorescent proteins across the cell over time, relative PKA activity, and changes therein, can be observed in subcellular space. FRET ratio images before and after treatment with either latrunculin A (100nM) or cytochalasin D (50nM) were compared to assess the effect of disrupted actin dynamics on PKA activity. To specifically monitor PKA activity in the leading edge, the leading edge was defined as the largest, dynamic protrusive structure using raw FRET images. Then, upon calculation of FRET ratio values, peak PKA activity was defined as FRET ratio values within the defined leading edge exceeding the cellular mean by 1.5 standard deviations. As expected, leading edge dynamics were rapidly inhibited (within 1 min) after addition of either drug (Supplemental Videos 1 and 2). However, neither actin polymerization inhibitor led to a significant decrease in the mass of relative leading edge PKA activity within 10 minutes of treatment (Supplemental Figure 1; Supplemental Videos 1 and 2). These observations, along with a previous study which showed that leading edge PKA activity in α4 integrin-expressing CHO cells gradually decreased 9-20 minutes after latrunculin A treatment [7], are in contrast to the rapid (within 2 min) and complete inhibition of PKA upon loss of contractility after treatment with blebbistatin and the even more rapid (within 30 sec) activation of PKA upon cell stretch [11]. Taken together, these data suggest that leading-edge PKA activity is not immediately or directly regulated by actin dynamics *per se* and suggest that PKA is coupled to changes in actomyosin contractility by means other than through actin polymerization.

### Disruption of cellular traction forces rapidly and transiently inhibits FAK activity within focal adhesions

The rapid effect of cell-ECM traction on PKA activity led us to consider other adhesion- and/or migration-related signaling pathways that are known to be mechanosensitive. Focal adhesion kinase (FAK) is a centrally important mediator of integrin-mediated adhesive signaling, regulator of cell migration, and component of mechanotransduction pathways [18, 19, 22, 23]. Thus, given that PKA activity is stimulated by integrin-mediated adhesion [29, 40], important for efficient cell migration [1, 3, 5, 9], and regulated by mechanical signaling [3, 41], FAK is well-positioned to be a regulator of leading edge PKA activity. Moreover, like PKA activity, FAK activity (as measured by autophosphorylation on Y397) also decreases upon inhibition of actomyosin contractility [42, 43]. Importantly, however, the effect of loss of contractility on FAK activity and/or autophosphorylation has, to date, only been observed after a period of 30-60 minutes, at which point focal adhesion morphology is dramatically altered [41-43]. In contrast, PKA activity decreases much faster after loss of tension - within 2 minutes - rapidly following the loss of cellular traction force and significantly preceding the structural dissolution of focal adhesions [11]. We therefore investigated whether there was an effect of blebbistatin on FAK activity at early timepoints that might correspond to the effect on PKA activity. SKOV-3 cells stimulated to migrate with 12.5ng/mL EGF, as in imaging experiments were treated with blebbistatin to assess the effect on FAK activity. Specifically, these migrating SKOV-3 cells were left untreated or were treated with blebbistatin for varying amounts of time (from 2 to 60 min) then harvested for immunoblotting or processed for immunofluorescence with antibodies against pY397 FAK (the autophosphorylation site). In agreement with published observations [42, 43], inhbition of contractility led to a decrease in total cellular FAK autophosphorylation by 30 min but had no apparent effect on autophosphorylation before 5 min (Figure 1A). However, when the phospho-FAK content within focal adhesions was assessed by immunofluorescence, as described in Figure 1B, there was a significant decrease in pY397-FAK (normalized to total focal adhesion area demarcated by vinculin staining as described in Figure 1B) within 2 minutes of blebbistatin treatment, (Figure 1C and D). This rapid decrease in FAK autophosphorylation was paralleled by a similar decrease in phosphorylation of paxillin, a major substrate of FAK, and Y118 (Figure 1C and E). These data suggest that a pool of active FAK is rapidly dephosphorylated within focal adhesions before the observable dissolution of the focal adhesion structures themselves. To our knowledge, this is the most immediate effect of myosin II inhibition on FAK phosphorylation described to date. Interestingly, the initial blebbistatin-mediated decrease in both pY397 FAK and pY118 paxillin levels was followed by a recovery by 5 min and a slight but steady and significant increase at later time points, becoming even higher than control levels as the focal adhesions dissolved and diminished in size (Figure 1C-E). This recovery in pY397-FAK levels is consistent with earlier observations that while focal adhesion size is greatly reduced 30 minutes after blebbistatin treatment, the level of phosphorylated FAK in remaining adhesive structures is comparable to the level found in mature focal adhesions in untreated cells [41]. This recovery notwithstanding, these observations confirm that FAK can be regulated by actomyosin contractility and establish that at least a portion of FAK activity responds with the same rapid kinetics as the loss of PKA activity to the loss of contractility.

**Figure 1:**
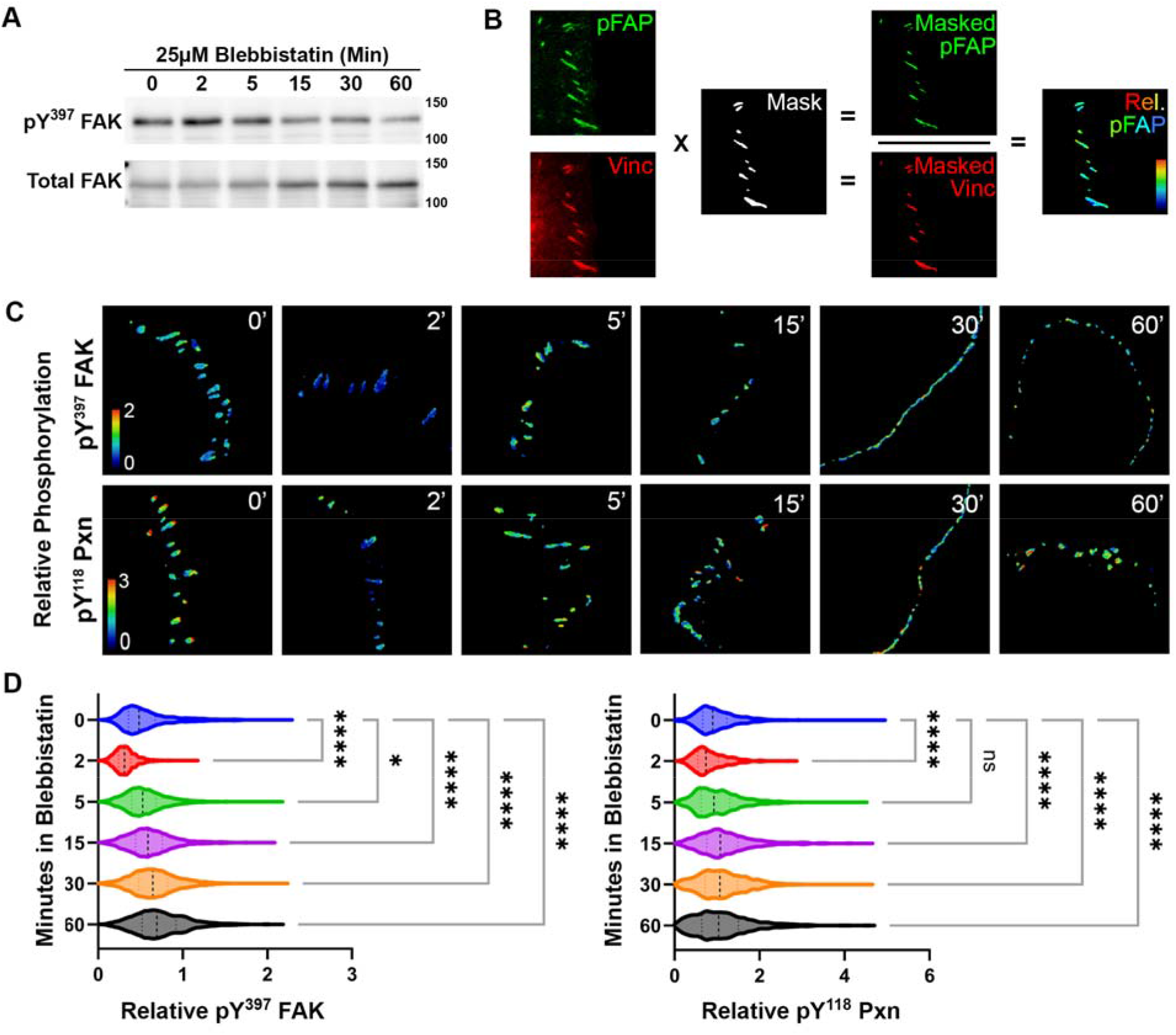
Blebbistatin rapidly but transiently inhibits the focal adhesion pool of FAK. (A) Western blot of whole cell lysates from cells treated with 25μM blebbistatin for 0, 2, 5, 15, 30, or 60 minutes probed for pY397 FAK or total FAK. (B) Schematic describing the method used to calculate and mask relative phospho-FAK and phospho-Pxn (<u>p</u>hosphorylated <u>F</u>ocal <u>A</u>dhesion <u>P</u>rotein (pFAP)) within focal adhesions. Briefly, SKOV-3 cells treated with 25 μM blebbistatin for 0, 2, 5, 15, 30, or 60 minutes were probed for vinculin, to demarcate total focal adhesions, and either pY397 FAK or pY118 Pxn (pFAP). pFAP signal was then divided by vinculin signal to give relative phosphoprotein images as shown in C. (C) Pseudo-colored relative phosphoprotein images for pY397 FAK (top) or pY118 Pxn (bottom) upon treatment with 25μM Blebbistatin for the indicated time period. (D) Relative phosphoprotein signal in each masked adhesion for each pY397 FAK (left) or pY118 Pxn (right). Kruskal-Wallis * p≤0.05, **** p≤0.0001.

### Pharmacologic inhibition of FAK decreases leading edge PKA activity

To directly test whether FAK activity was required for leading edge PKA activity, SKOV-3 cells transiently expressing pmAKAR3 were analyzed by FRET microscopy before and after treatment with PF-562271 (PF271), a potent, ATP-competitive FAK inhibitor. Treatment with 250nM PF271 led to a rapid decrease in membrane-proximal leading edge PKA activity within 10 minutes of treatment (Figure 2A-D; Supplemental Video 3). This effect did not change the ability of PKA to respond to increases in cAMP using a saturating dose of forskolin (to activate adenylyl cyclases) and IBMX (to inhibit phosphodiesterases; Supplemental Figure 2), though the measurement of this effect was limited by the dynamic range of the biosensor. Importantly, the concentration used here was 2.5-fold lower than the reported *in vitro* IC_50_ of PF271 against PKA directly [44]. Nonetheless, to ensure that the decrease in PKA activity was not an off-target effect of PF271, the assay was repeated with two additional FAK inhibitors - PND1186 and the lesser known, but structurally dissimilar, FAK inhibitor 14 (also known as Y15). Treatment with FAK inhibitor 14 caused a rapid and complete reduction of leading edge PKA activity, even more striking than that caused by PF271 (Figure 2D). While we found no data reporting any direct effect of FAK inhibitor 14 on PKA activity, this inhibitor is reported as highly selective for FAK activity, having no measurable effect on even highly similar tyrosine kinases such as Pyk2 or Src [45-47]. Treatment with a third FAK inhibitor, PND1186 (also called VS-4718), led to a distinct effect. Whereas peaks of PKA activity in the leading edge were diminished, as seen with the previous inhibitors, overall PKA activity in other parts of the cell rose (Supplemental Figure 3 A and B). While the reason for this increase is unknown, it might be explained by *in vitro* studies that showed PND1186 can cause a slight direct increase in PKA activity [48]. Taken together, the demonstration that three chemically distinct inhibitors of FAK rapidly decreased leading edge PKA activity solidly suggests a functional connection between these two pathways.

**Figure 2:**
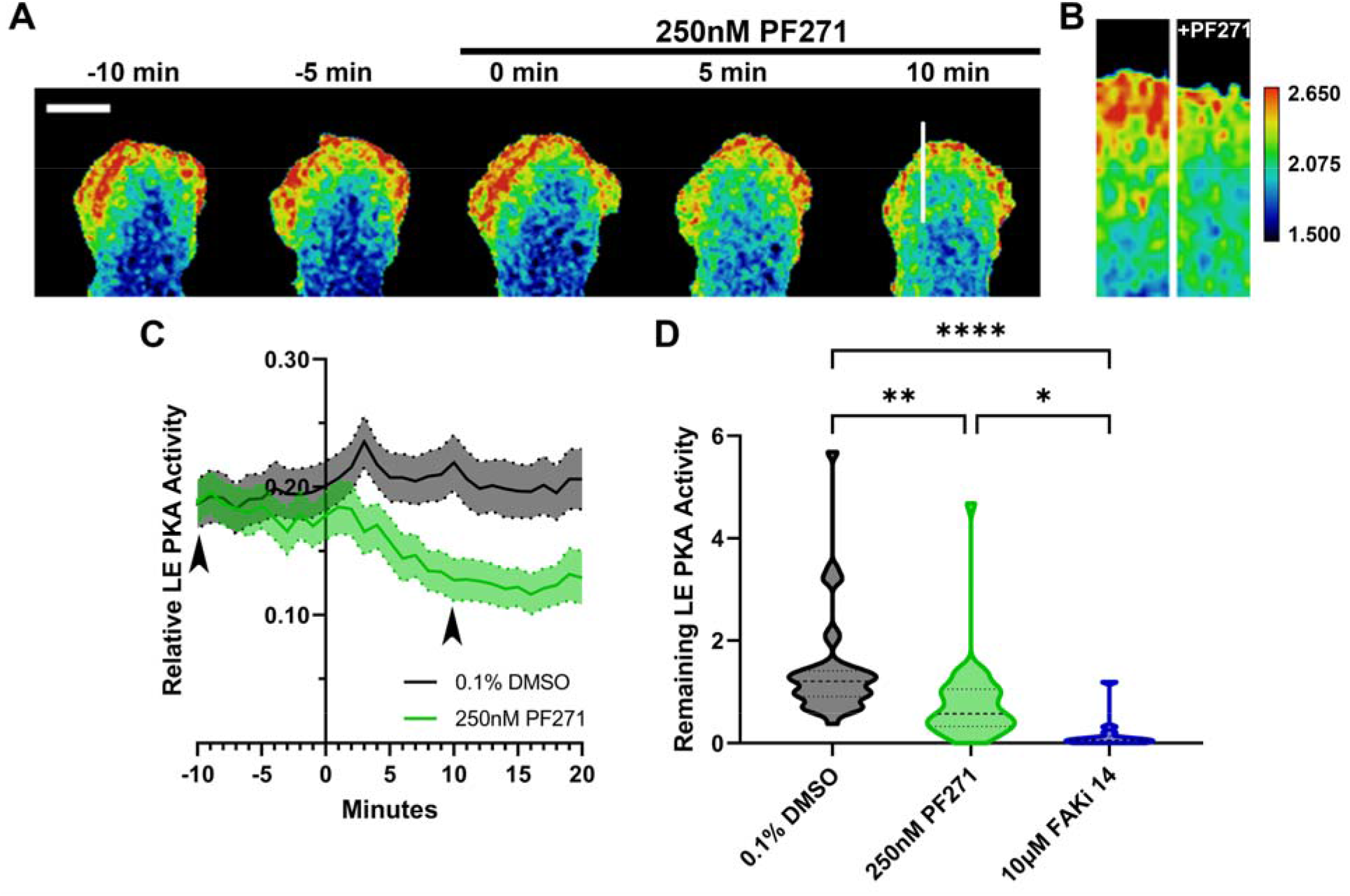
Inhibiting FAK activity rapidly decreases leading edge PKA activity. (A) FRET ratio image montage of the leading edge of an SKOV-3 cell expressing pm-AKAR3 treated with 250nM PF271 at time 0, scale=10μm. (B) Kymograph of leading edge PKA activity through line in panel A. (C) Relative peak PKA activity in the leading edge of cells over time treated with vehicle control, 0.1% DMSO (N=3 replicates, n=36 migrating cells), and cells treated with 250nM PF271 (N=3 replicates, n=36 migrating cells). Arrowheads indicate timepoints used to calculate pharmacological effect in D. (D) Peak PKA activity remaining in the leading edge 10 minutes after treatment compared to that 10 minutes before treatment with 0.1% DMSO, 250nM PF271, and 10μM FAKi 14 (N=1 replicates, n=10 migrating cells). Kruskal-Wallis with multiple comparisons performed for all pharmacologic treatments (see also Supplemental Figures 1 and 3), * p≤0.05, ** p≤0.01, **** p<0.0001. Comparisons shown are a subset of all comparisons assessed by Kruskal-Wallis test. Comprehensive graph including test results for all pharmacologic treatments shown in Supplemental Figure 4.

**Figure 3:**
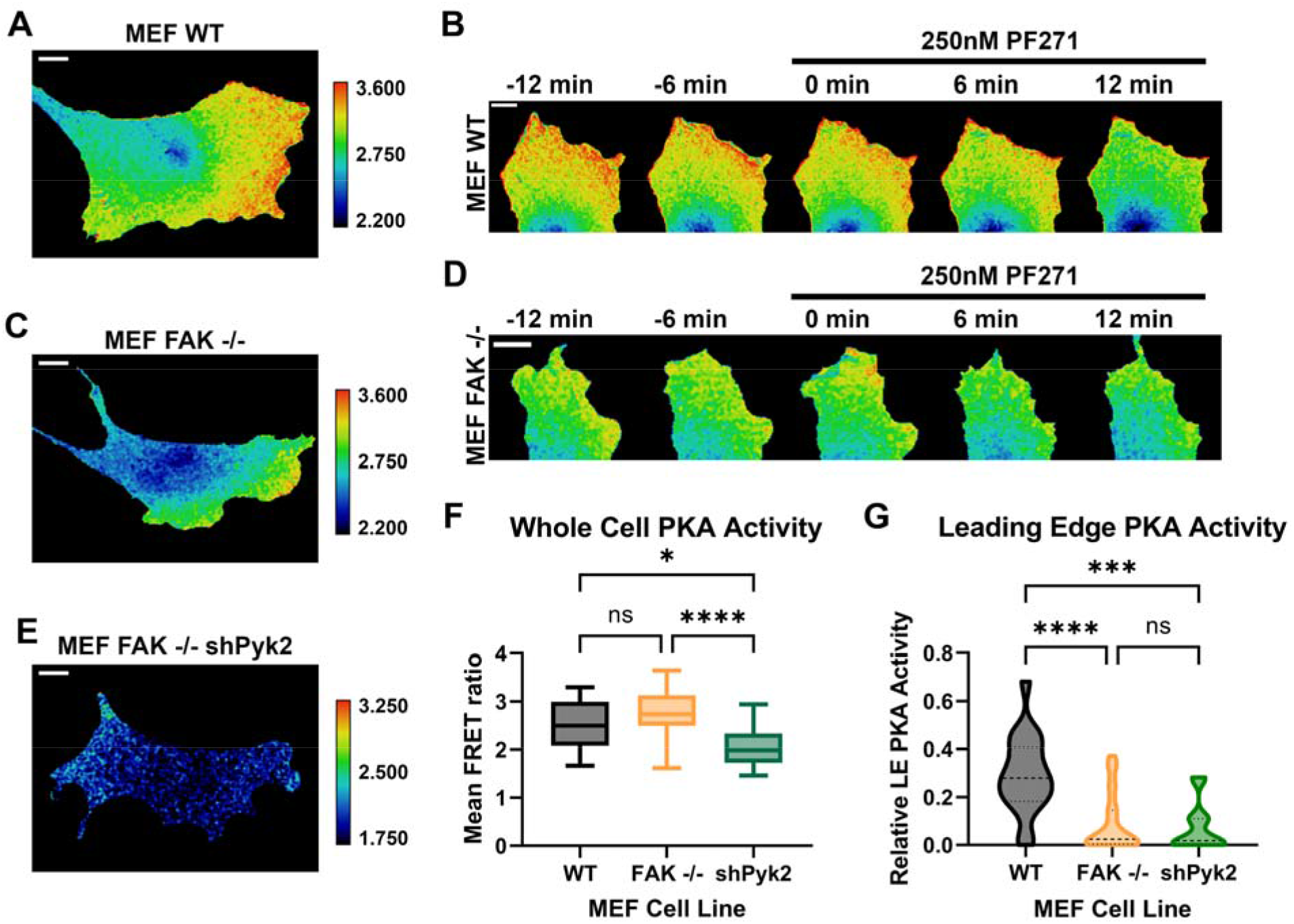
Fibroblasts lacking FAK exhibit decreased leading edge PKA activity. A) FRET ratio image of a migrating wildtype MEF cell expressing pm-AKAR3, PKA activity reporter, scale bar=10μm. (B) Montage of FRET ratio in the leading edge of a wildtype MEF cell upon treatment with 250nM PF271 showing decreased leading edge PKA activity six and twelve minutes after treatment, scale bar=10μm. (C) FRET ratio image of a migrating FAK-null MEF cell expressing pm-AKAR3, scale bar=10μm. Note the uniform LUT scaling in panels A and C. (D) Montage of FRET ratio in the leading edge of a FAK-null MEF cell upon treatment with 250nM PF271, showing very slightly decreased leading edge PKA activity six and twelve minutes after treatment, scale bar=10μm. (E) FRET ratio image of a MEF FAK-null cell stably expressing shPyk2 and transiently expressing pm-AKAR3, scale bar=10μm. LUT scaling enhanced due to decreased mean FRET ratios in shPyk2 cells. (F) Whole cell PKA activity as assessed by mean FRET ratio values for entire cell in wildtype MEF (N=5 replicates, n=30 migrating cells), FAK-null MEF (N=3 replicates, n=27 migrating cells), and FAK-null/shPyk2 MEF (N=5 replicates, n=24 migrating cells) cells. Comparisons shown are a subset of all comparisons assessed by Kruskal-Wallis test. Comprehensive graph including all test results found in Supplemental Figure 5A. (G) Relative leading edge PKA activity in wildtype MEF (N=5 replicates, n=19 migrating cells), FAK-null MEF (N=3 replicates, n=26 migrating cells), and FAK-null/shPyk2 MEF (N=4 replicates, n=14 migrating cells) cells. Comparisons shown are a subset of all comparisons assessed by Kruskal-Wallis test. Comprehensive graph including all test results found in Supplemental Figure 5B. * p≤0.05, *** p≤0.001, **** p≤0.0001.

### Genetic loss of FAK leads to decreased PKA activity in the leading edge

Despite both the current observations as well as the parallels between FAK and PKA in terms of their regulation by and role in adhesion, migration, and mechanotransduction, there are no reports of FAK acting upstream of PKA activity and no obvious mechanism or target through which FAK might exert its effect. Thus, to eliminate the possibility of off-target effects of pharmacologic inhibition and to further confirm this novel regulatory connection, leading edge PKA activity was observed in wild-type mouse embryonic fibroblasts (MEFs) as well as MEFs isolated from FAK-null embryos [49]. These cell lines were chosen given their extensive characterization and validation in previous studies [48 and many more]. Similar to SKOV-3 and other cell types previously assessed for PKA activity during migration [1, 3, 5, 7-11, 50], wild-type MEF cells exhibit leading edge PKA events (Figure 3A; Supplemental Video 4). Also, as in SKOV-3 cells, this localized PKA activity is rapidly decreased upon pharmacologic inhibition of FAK by PF271 (Figure 3B; Supplemental Video 4). However, FAK-null MEFs showed significantly lower peak and average PKA activity in the leading edge compared to wild-type MEFs (Figure 3C, D, and G; Supplemental Video 5). Previous characterization of FAK-null MEFs have revealed partial migration defects in these cell lines, and while their speed and persistence are altered, cells still develop protrusive leading edges and are able to translocate [CITE] and thus, definition of the leading edge was possible in these cells. Of note, the whole-cell mean FRET ratios of the FAK-null cells was not significantly different than that of wild-type MEFs and a minimal level of membrane-proximal PKA activity was retained in protrusive regions (Figure 3C, F, and G).

Interestingly, the low level of leading edge PKA activity observed in FAK-null cells was further diminished by treatment with PF271 (Figure 3D; Supplemental Video 5), suggesting that FAK relative Pyk2 may be compensating for the loss of FAK in this system. Pyk2 is a related tyrosine kinase which shares 45% amino acid identity with FAK and is upregulated and highly active in FAK-null cells [51]. Prior studies have shown that this increase in Pyk2 abundance and activity can partially compensate for the loss of FAK in some contexts [51, 52]. Further, Pyk2 is also inhibited by PF271, albeit with a higher IC_50_ than FAK [44]. Thus, we assessed whether loss of Pyk2 would further diminish PKA activity in the FAK-null MEFs. To this end, we assessed PKA activity in FAK-null MEFs stably expressing a short hairpin RNA against Pyk2 to knock down Pyk2 activity (shPyk2). Total PKA activity in the FAK-null/shPyk2 cells was significantly diminished compared to both the wild-type and FAK-null cells, as measured by whole cell FRET ratio (Figure 3E and F; Supplemental Video 6). Also, though the relative mass of leading edge PKA activity in FAK-null/shPyk2 cells was lower than in FAK-null cells, this decrease was not statistically significant, likely due to the already low level of activity in the FAK-nulls and the limited number of FAK-null/shPyk2 cells per experiment that were amenable to analysis due to low transfection efficiency (Figure 3E and G). Taken together, these genetic data corroborate the observations from the pharmacological inhibitors and demonstrate that FAK - and to a lesser extent, Pyk2 - is required for leading edge PKA activity in migrating cells.

### Leading edge PKA activity in FAK-null cells is rescued by wild-type but not kinase-dead FAK

In addition to its tyrosine kinase activity, FAK has been shown to exhibit important cellular effects by acting as a scaffold that physically organizes various signaling intermediates [53]. To further corroborate the pharmacologic inhibitor data and confirm that FAK kinase activity rather its structural and scaffolding functions were required for regulation of leading edge PKA activity, FAK-null MEFs were co-transfected with constructs expressing pm-AKAR3 and either mCherry-tagged wild-type or kinase-dead (R454) FAK, or mCherry-paxillin as a control. Expression of target proteins was confirmed by assessing mCherry fluorescence. Re-expression of wild-type FAK rescued leading edge PKA activity to the level of wildtype cells (Figure 4A and E; Supplemental Video 7). As expected, expression of mCherry-paxillin was unable to rescue leading edge PKA activity beyond that of the parent FAK-null cell line (Figure 4 B and E; Supplemental Video 8). Similarly, cells expressing the kinase dead mCherry-R454 FAK exhibited limited PKA activity in the leading edge, similar to FAK-null cells alone or those expressing mCherry-paxillin (Figure 4 C and E; Supplemental Video 9). These data validate the causal connection between FAK function and leading edge PKA activity and demonstrate that FAK kinase activity, and not its scaffolding or other functions, is required for its regulation of PKA activity in the leading edge of migrating cells.

**Figure 4:**
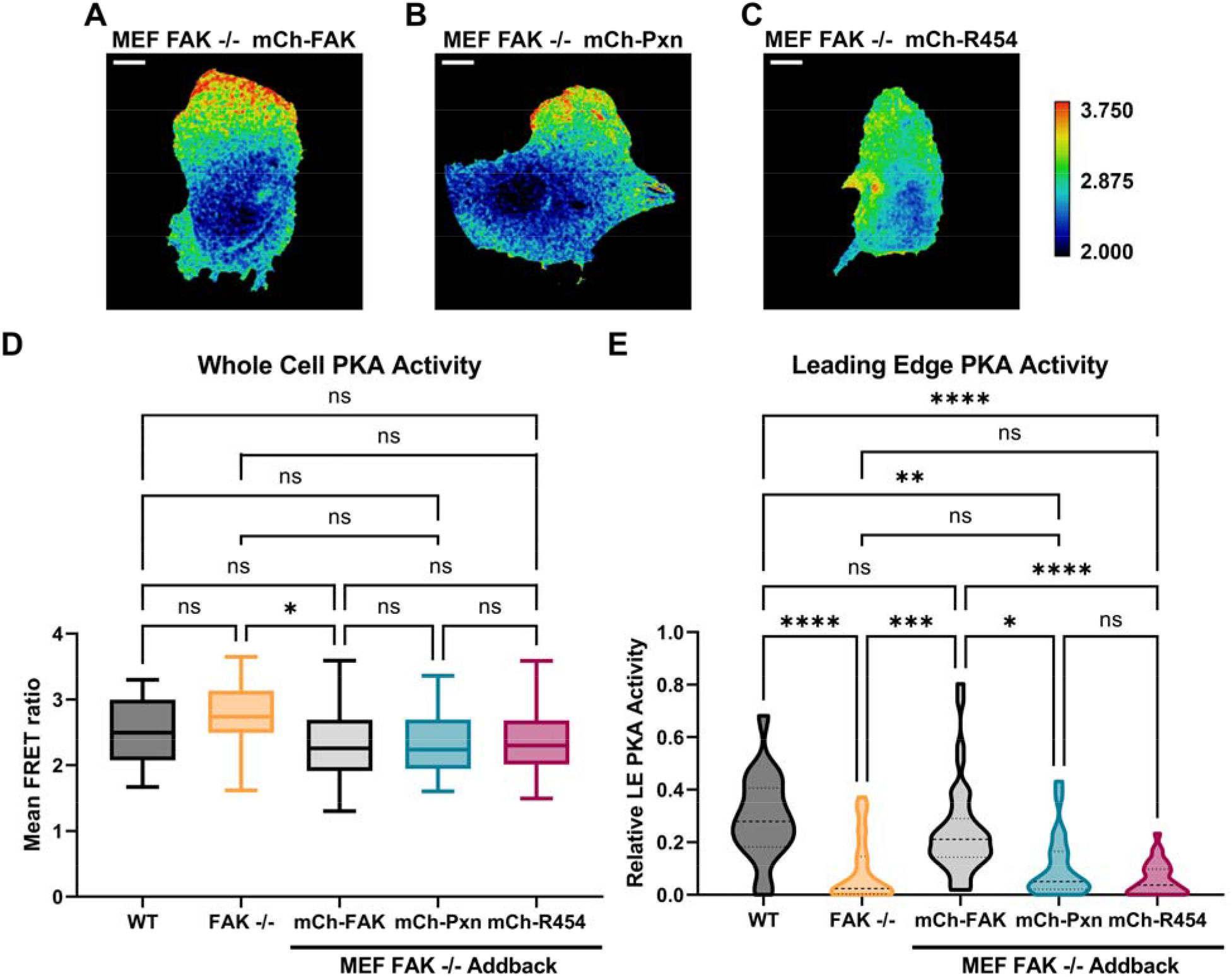
Leading edge PKA activity is rescued by wild-type but not kinase-dead FAK. (A) FRET ratio image of a migrating FAK-null MEF cell expressing mCherry-(mCh-)FAK and pm-AKAR3, scale bar=10μm. (B) FRET ratio image of a migrating FAK-null MEF cell expressing mCh-paxillin and pm-AKAR3, scale bar=10μm. (C) FRET ratio image of a MEF FAK-null cell expressing mCh-R454 FAK and pm-AKAR3, scale bar=10μm. Note the LUT scaling is the same for A-C. (D) Whole cell PKA activity as assessed by mean FRET ratio values for entire cell. mCh-FAK (N=3 replicates, n=42 migrating cells), mCh-Paxillin (N=3 replicates, n=22 migrating cells), or mCh-R454 FAK (N=3 replicates, n=30 migrating cells). Comparisons shown are a subset of all comparisons assessed by Kruskal-Wallis test. Comprehensive graph including all test results found in Supplemental Figure 5A. (E) Relative leading edge PKA activity in FAK-null MEF cells expressing mCh-FAK (N=3 replicates, n=42 migrating cells), mCh-Paxillin (N=3 replicates, n=22 migrating cells), or mCh-R454 FAK (N=3 replicates, n=30 migrating cells). Comparisons shown are a subset of all comparisons assessed by Kruskal-Wallis test. Comprehensive graph including all test results found in Supplemental Figure 5B. * p≤0.05, ** p≤0.01, *** p≤0.001, **** p≤0.0001.

## DISCUSSION

We have identified a novel relationship whereby FAK promotes PKA activity in the leading edge of migrating SKOV-3 ovarian cancer cells and MEF cells. This relationship is sensitive to both pharmacologic and genetic inhibition of FAK and is dependent on FAK kinase activity. It is rather surprising that FAK inhibition has such a marked effect on PKA activity in the leading edge. The canonical pathway leading to activation of PKA, the generation of cAMP downstream of adenylyl cyclases (ACs), heterotrimeric G-proteins, and GPCRs, is well characterized and ostensibly wholly independent of tyrosine phosphorylation. There is no precedent suggesting the mechanism by which this regulation occurs. It is possible that FAK is acting on regulatory targets upstream of PKA, such as heterotrimeric G proteins, ACs, or phosphodiesterases, that control the local availability of cAMP. To our knowledge, however, there are no published reports of FAK activity contributing to the generation of cAMP, in any system. There is evidence to support the regulation of FAK downstream of ACs, both promoting FAK activity downstream of increased AC activity [31] or norepinephrine stimulation [54], and decreasing FAK activity after stimulation of ACs in microdomains [55]. Further, inhibition of PKA has been shown to delay dephosphorylation of FAK upon cellular detachment, putting PKA upstream of FAK in anchorage-dependent signaling [29]. Yet no mechanism for these relationships has been identified, nor has the reciprocal relationship, whereby FAK modulates the activity of adenylyl cyclases, been demonstrated.

While other tyrosine kinases such as EGFR and Fyn have been shown to directly phosphorylate the PKA catalytic domain [35, 36], these modifications lead to only a modest increase in the catalytic efficiency of PKA *in vitro*, and the direct PKA phosphorylation by FAK has not been investigated. Thus, while phosphorylation of PKA subunits by FAK has not been shown, it is possible that FAK, like EGFR and Fyn, may directly modify and thus regulate PKA catalytic subunits. However, if FAK were directly phosphorylating PKA, it seems unlikely that we would see a marked decrease in PKA activity within minutes of FAK inhibition as we see in Figure 2. If, similar to EGFR and Fyn, phosphorylation of PKA by FAK slightly increased PKA catalytic efficiency, then inhibiting FAK would cease ongoing phosphorylation of PKA. A decrease in PKA activity would then rely on swift, overwhelming action by tyrosine phosphatases, which would likely only modestly decrease the catalytic response of PKA to the unchanged availability of cAMP. Therefore, it is more likely that FAK is acting upstream, potentiating PKA through increased availability of cAMP, allowing for more immediate and marked change through signal amplification via second messenger production.

Moreover, using the R454 kinase-dead FAK construct in FAK-null cells, we have demonstrated that it is FAK kinase activity, and not one of the numerous scaffolding functions of FAK, that is responsible for regulation of PKA. FAK is a large protein known for its ability to scaffold higher order signaling complexes through its binding with numerous other proteins—other kinases such as Src, adapter molecules such as p130Cas, and focal adhesion components like talin and paxillin [53, 56-59]. Our results suggest that FAK is acting to influence PKA activity more directly than by scaffolding some components of the signaling pathway in a way that requires the input of its kinase activity.

With these results, it is unclear whether mechanical inputs, PKA, and FAK exist in a linear hierarchy where FAK acts as an intermediate signaling node. Prior experiments suggested that FAK inhibition does not rapidly change actomyosin contractility [41]. If FAK is acting to transmit mechanical signals to PKA, we would expect that disrupting contractility would lead to rapid decreases in FAK activity. Though we have shown that at least some pools of FAK are rapidly sensitive to contractile forces (within 2 minutes), the data presented do not categorically place FAK activity in the same pathway as mechanical regulation of PKA. While the loss of contractility rapidly inhibited FAK activity in focal adhesions, our prior work has shown that leading edge PKA events are not directly spatially related to focal adhesions [11]. Additionally, FAK functions in distinct pools within the cell, not always localized to focal adhesions [60]. Therefore, further work must be done at higher time resolution to identify whether a subcellular pool of FAK activity is contributing to PKA activity in migrating cells. This can be addressed using subcellular fractionation techniques to sensitize western blotting to small, subcellular changes in FAK activity downstream of myosin II inhibition.

The novel involvement of FAK in leading edge PKA activity is intriguing regardless of whether FAK is involved specifically in its mechanical regulation. PKA is active in the leading edge of numerous cell types, is required for the migration of many different types of cells, and has important downstream targets involved in many aspects of this complex process [3]. FAK is similarly widely required for cell migration, adhesion, and mechanotransduction, yet these two principal kinases are canonically regarded to function separately. The identification of crosstalk between FAK and PKA represents an important finding in the regulation of migration and implies that these ubiquitous but heretofore disparate pathways may crosstalk in other cellular processes. This represents a wholly new aspect of regulation of the PKA pathway and an itinerant new avenue of investigation to identify its extent and underlying mechanism.

## MATERIALS AND METHODS

### Reagents and Cell Culture

SKOV-3 cells, purchased from the American Type Culture Collection, were maintained in DMEM supplemented with 10% fetal bovine serum below 60% confluency. Cells plated for transfection were allowed to reach 70% confluency and transfected with FuGene 6 (Promega) according to manufacturer’s protocol. After 30 hours, transfected SKOV-3 cells were trypsinized and cryopreserved at -80°C in DMEM 10% fetal bovine serum, 10% DMSO.

MEF WT, FAK KO, and FAK KO shPyk2 cells were obtained from David Schlaepfer and maintained in DMEM supplemented with 10% fetal bovine serum on dishes coated with 0.1% gelatin [49]. Cells were transfected at 70% confluency, replated at 24 hours, and cryopreserved after 48 hours at -80°C in DMEM 10% fetal bovine serum, 10% DMSO.

Blebbistatin was purchased from Tocris. Acrylamide and N,NI-methylenebisacrylamide were purchased from National Diagnostics. Tetramethylethylenediamine (TEMED), ammonium persulfate (APS), and bovine serum albumin (BSA), along with other sundry chemicals, were purchased from Sigma (St. Louis, MO). Plasmids used in this work included PKA biosensor pmAKAR3 (Allen and Zhang, 2006) from Jin Zhang (Johns Hopkins University pmCherry-FAK from Addgene (plasmid #35039).

### FRET Microscopy

Imaging dishes (29mm; CellVis) were coated overnight at 4°C with 10μg/ml fibronectin. SKOV-3 cells were thawed and plated at 25,000 cells per dish in DMEM supplemented with 10% fetal bovine serum. Four hours after plating, cells were rinsed and media was exchanged to Ringer’s Solution supplemented with 12.5ng/ml epidermal growth factor. Cells were allowed to equilibrate for at least 45 min before imaging. Imaging began 20 minutes before drug treatment and extended 20 minutes beyond treatment to assess the effect of each drug.

MEF cells were thawed and plated at 20,000 cells per dish in DMEM supplemented with 10% fetal bovine serum. Eighteen hours after plating, cells were rinsed and media was exchanged with Ringer’s Solution supplemented with 10ng/ml plasma derived growth factor-BB.

CFP and FRET images were acquired simultaneously using a Nikon TiE inverted fluorescent microscope and the Cairn OptoSplitII optical splitter. Temperature was maintained at 37°C. FRET images were acquired using 2×2 binning in Nikon Elements imaging software.

### FRET Image Analysis

FRET images were calculated using a protocol based on Broussard *et al*. as described previously [11, 61]. Raw CFP and FRET images were prepared using median filter, radius1 and background was subtracted from each channel using the mean background pixel value for each image. The FRET image was then corrected for CFP bleed-through and division of the corrected FRET image by the CFP image resulted in the FRET ratio image.

FRET ratio images were then masked as follows. The corrected FRET and CFP images were added together and the resulting image was sharpened using an unsharp mask, radius 10. This image was turned into a binary mask via thresholding. The ratio image was then multiplied by this binary mask, turning background pixels to NaN.

### Quantification of Relative Leading Edge PKA Activity

As previously reported, membrane-proximal PKA activity is increased in the leading edge of migrating cells as compared to the rest of the cell. Therefore, in order to quantify this elevated leading edge activity and observe it over the course of an experiment, the area of increased signal in the leading edge, relative to the size of the leading edge, was calculated for each frame of the movie. The leading edge of each moving cell was captured manually in a region of interest. The leading edge of the cell was defined as the largest dynamic protrusion using the raw captured FRET image, prior to the calculation of FRET ratio images. Given the protruding nature of the leading edge, this portion of the cell was registered over the length of the experiment to spatially isolate the leading edge for further calculation of signal mass. The amount of relative leading edge activity was calculated as follows. The mean FRET ratio value of the entire migrating cell was calculated over all frames captured before pharmacologic manipulation. Next, the elevated PKA activity mass in each frame was defined as the number of pixels in the leading edge whose FRET ratio value measured more than 1.5 standard deviations above the mean pixel value, defined above. To calculate relative leading edge PKA activity, this signal mass was divided by the total number of pixels in the leading edge. The leading edge activity remaining after a given treatment was calculated by dividing the relative LE PKA activity 10 minutes after by that 10 minutes before treatment. For MEF cell lines, whole cell mean FRET ratio values and relative leading edge activity values werecalculated by averaging the relative leading edge activity over 21 minutes, or 7 frames, of the movie, absent of any treatment, while the cell migrated.

### Statistical Analysis

In all cases, values were bounded asymmetrically by zero. Therefore, a nonparametric ANOVA was used to analyze each data set. A single Kruskal-Wallis comparison was performed on all pharmacologic treatments of SKOV3 cells. The results of this comparison are discussed throughout several sections of the results and are thus presented in several figures. Comprehensive results of the Kruskal-Wallis comparison of every pharmacologic intervention performed are shown in Supplemental Figure 4. Subsets of this comparison are presented in Figure 2 and in Supplemental Figures 1 and 3 to highlight the effects of specific treatments. In each occurrence, the significance level shown is that relative to all treatments, as calculated in the comprehensive Kruskal-Wallis comparison. Similarly, a single Kruskal-Wallis comparison of all MEF cell lines appears in Supplemental Figure 5. These comparisons are shown in part in Figures 3 and 4.

### Immunofluorescence

SKOV-3 cells were plated in DMEM supplemented with 10% fetal bovine serum on coverslips coated with 10μg/mL fibronectin overnight then treated with 25 uM blebbistatin for the indicated periods. Cells were then fixed in 4% PFA in PBS for 15 min at RT, permeabilized in 0.25% triton X100 in PBS for 10 min at RT with gentle shaking, and blocked for at least 1 h in 4% low-IgG BSA + 0.1% tween-20 in PBS. Coverslips were stained for 1 h at RT with antibodies against vinculin (hVin1, Sigma; 1:400) and either (phosphoTyr397-FAK (Cat# 44-625G, ThermoFisher; 1:50) or phosphoTyr118-paxillin (Cat# 44-722G; ThermoFisher; 1:100) diluted in blocking solution. Coverslips were washed 4 times for 7 min in PBS, then incubated in secondary antibodies (Alexa594-ocnjugated donkey anti-mouse and Alexa488-conjugated anti-rabbit, both from Abcam and both diluted 1:250 in block) and washed again as above. Coverslips were mounted on slides using Prolong Glass Antifade (ThermoFisher) and imaged as described elsewhere [41].

### Calculation of Relative Phosphorylation of Focal Adhesion Proteins

Vinculin staining was used to represent total focal adhesion signal, both to demarcate the footprint of each focal adhesion by creating a mask for the phosphorylated focal adhesion protein signal (pFAP) and as a comparator for levels of FAK or paxillin phosphorylation. The steps of this calculation are laid out schematically in Figure 1B. Vinculin signal was used to create a binary focal adhesion mask which was multiplied by the pFAP signal. Masked vinculin immunofluorescent images were then used as a denominator for each masked pFAP image. Dividing masked pFAP images by masked vinculin images produced images depicting relative phosphorylation of the given pFAP which were pseudo-colored to highlight the varying level of relative phosphorylation of paxillin or FAK, as shown in Figure 1C,

### Western Blot

SKOV-3 cells were plated in DMEM supplemented with 10% fetal bovine serum on 60 mm tissue culture dishes previously coated in 10μg/mL fibronectin. Following a similar protocol to imaging parameters, at ∼70% confluency, cells were stimulated with 12.5ng/mL EGF for 30 minutes prior to drug treatments. Timed treatments with 25μM blebbistatin were ended by a double rinse in ice-cold complete PBS followed by harvest in lysis buffer containing 50mM Tris, pH 8.0, 150mM NaCl, 5mM EDTA, 5% glycerol, 1% Triton-X 100, 25mM NaF, 2mM Na_2_VO_4_ supplemented with 1x protease inhibitor cocktail (Thermo Scientific). Cells were freeze-thawed twice in liquid nitrogen and whole cell extracts were collected upon centrifugation at 16,000 x g. 10μg of protein from each treatment condition was separated by SDS-PAGE and transferred to PVDF membrane. Membranes were blocked with 5% nonfat dry milk and immunoblotted with antibodies against pY397-FAK (Invitrogen 44-624G) or total FAK (Millipore, 05-537).

## Supplemental Figure Legends

**Supplemental Figure 1:**
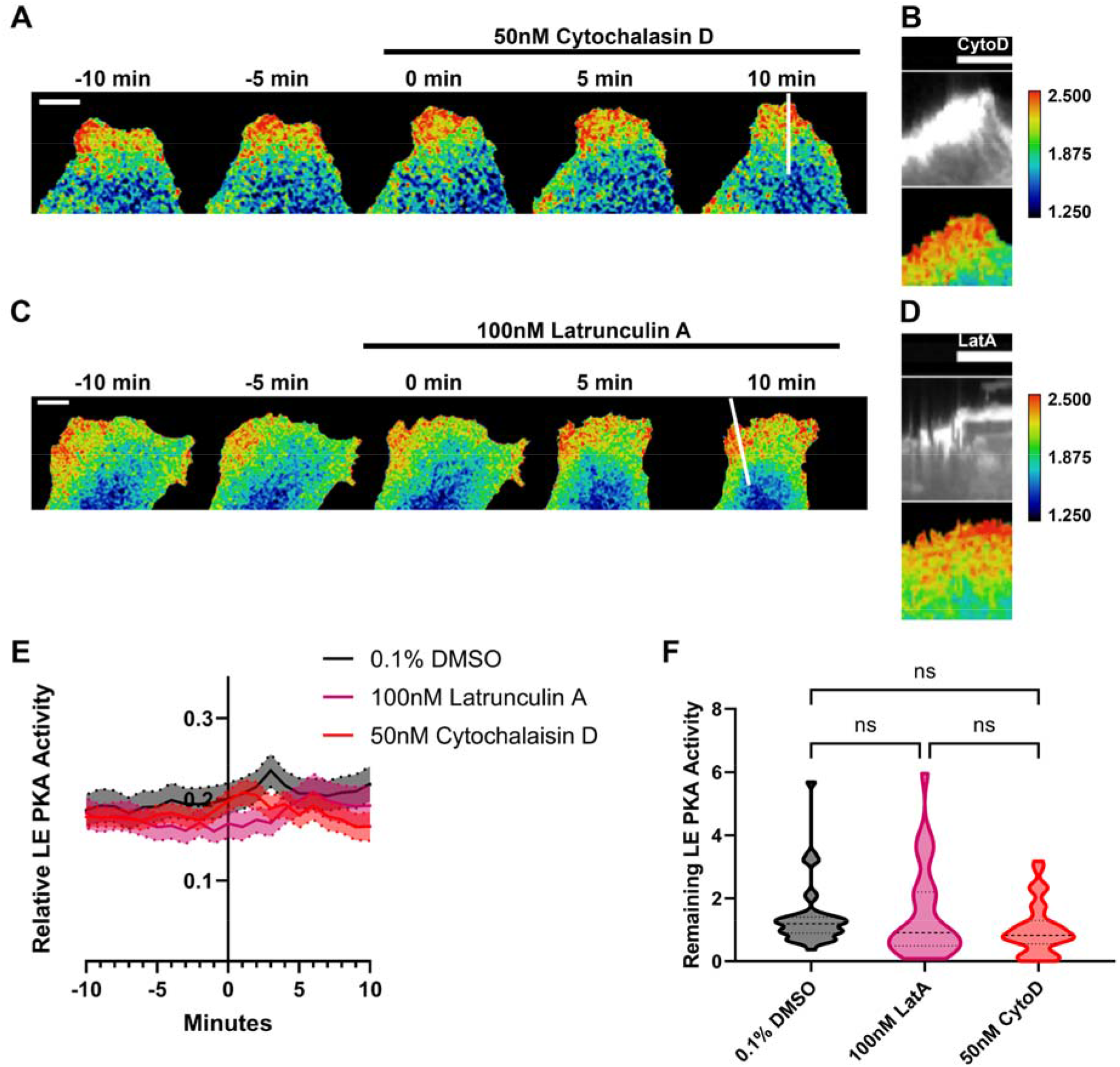
Inhibiting actin polymerization does not rapidly decrease leading edge PKA activity (A) FRET ratio image montage of the leading edge of an SKOV-3 cell expressing pm-AKAR3 treated with 50nM Cytochalasin D at time 0, scale=10μm. (B) Kymograph of pm-AKAR3 raw YFP emission (greyscale, intentionally over-brightened for ease of visualization; *top*) and leading edge PKA activity (pseudocolored; *bottom*) through the linear region of interest (ROI) in panel A. (C) FRET ratio image montage of the leading edge of an SKOV-3 cell expressing pm-AKAR3 treated with 100nM Latrunculin A at time 0. Scale=10μm (D) Kymograph of pm-AKAR3 raw YFP emission and leading edge PKA activity through the ROI in panel C. (E) Relative peak PKA activity in the leading edge of cells over time treated with vehicle control, 0.1% DMSO (N=3 replicates, n=36 migrating cells), and cells treated with 50nM Cytochalasin D (N=3 replicates, n=34 migrating cells) or 100nM Latrunculin A (N=3 replicates, n=28 migrating cells). (D) Peak PKA activity remaining in the leading edge 10 minutes after treatment compared to that 10 minutes before treatment with 0.1% DMSO, 50nM Cytochalasin D, or 100nM Latrunculin A. Kruskal-Wallis with multiple comparisons. Comparisons shown are a subset of all comparisons assessed by Kruskal-Wallis test. Comprehensive graph including all test results found in Supplemental Figure 4.

**Supplemental Figure 2:**
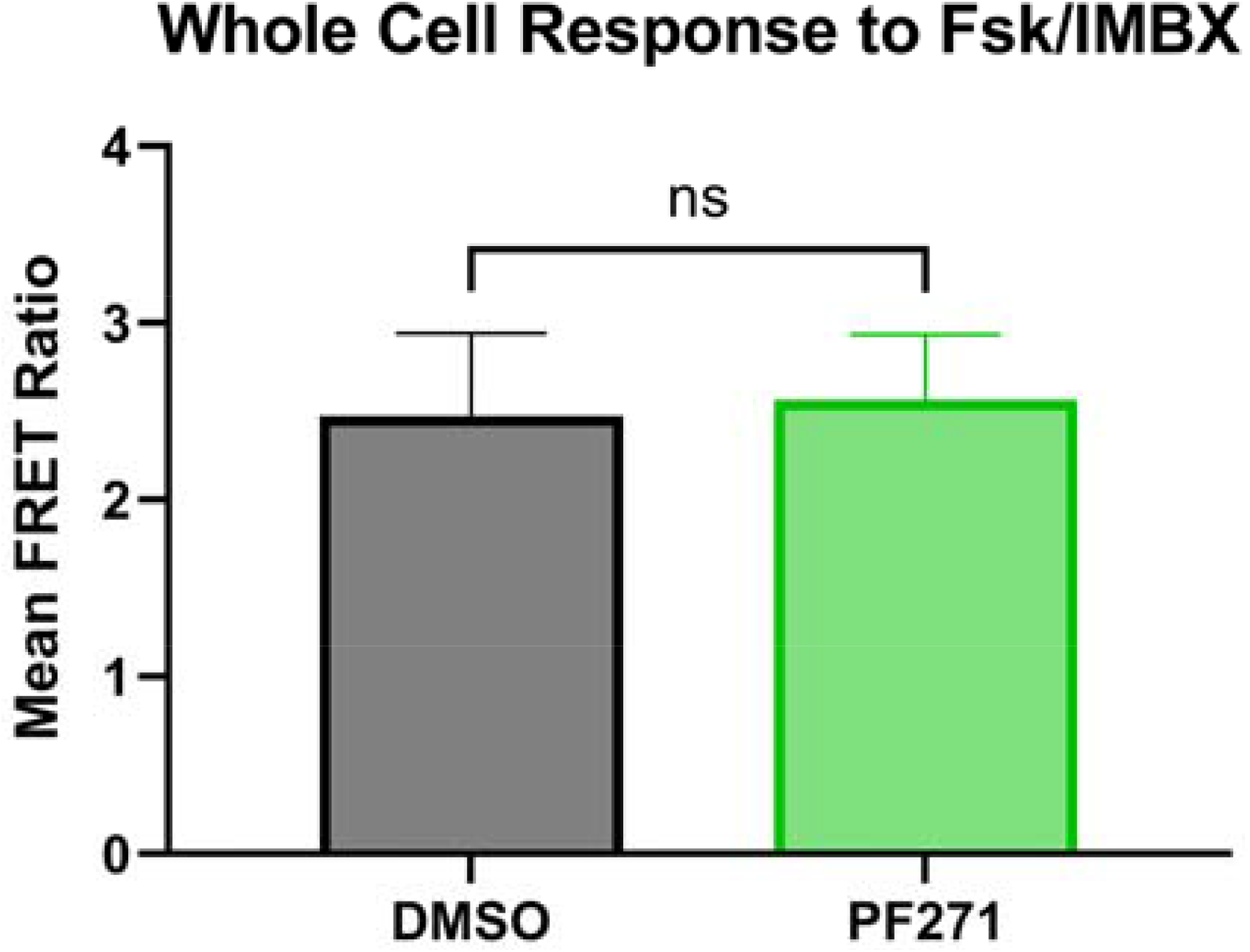
Whole cell FRET ratio to Forskolin and IBMX in SKOV-3 cells after treatment with PF271. After treatment with PF271 or DMSO control, cells from Figure 2 were treated with Forskolin and IBMX to increase intracellular cAMP and assess maximum PKA signal. No significant difference was found between the cell means.

**Supplemental Figure 3:**
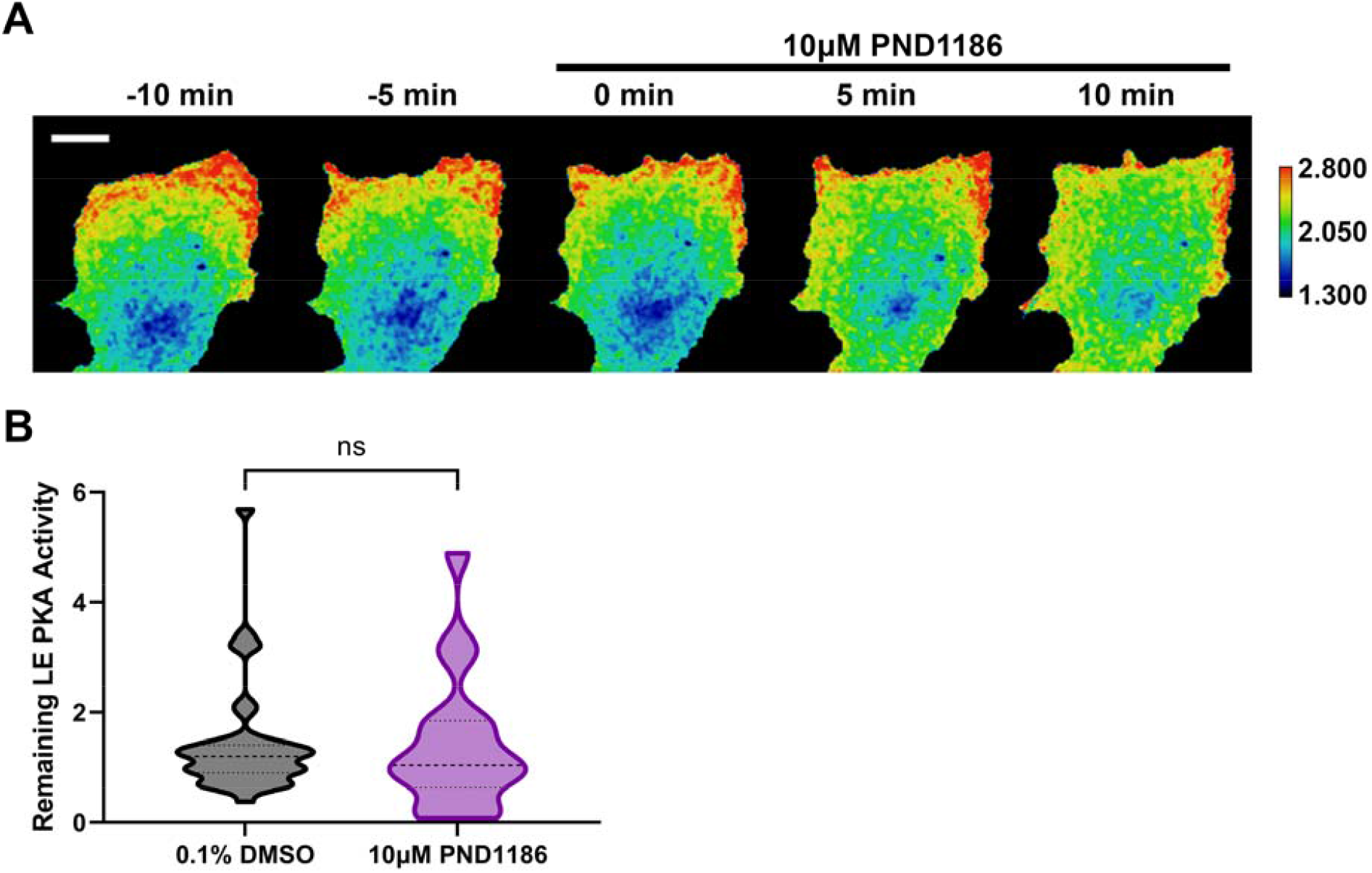
Inhibiting FAK with PND1186 leads to decrease in leading edge PKA activity, but slight whole cell increase. (A) FRET ratio image montage of the leading edge of an SKOV-3 cell expressing pm-AKAR3 treated with 10μM at time 0, scale=10μm (B) Peak PKA activity remaining in the leading edge 10 minutes after treatment compared to that 10 minutes before treatment with 0.1% DMSO (N=3 replicates, n=36 migrating cells) or 10μM PND1186 (N=2 replicates, n=18 migrating cells). Kruskal-Wallis with multiple comparisons. Comparisons shown are a subset of all comparisons assessed by Kruskal-Wallis test. Comprehensive graph including all test results found in Supplemental Figure 4.

**Supplemental Figure 4:**
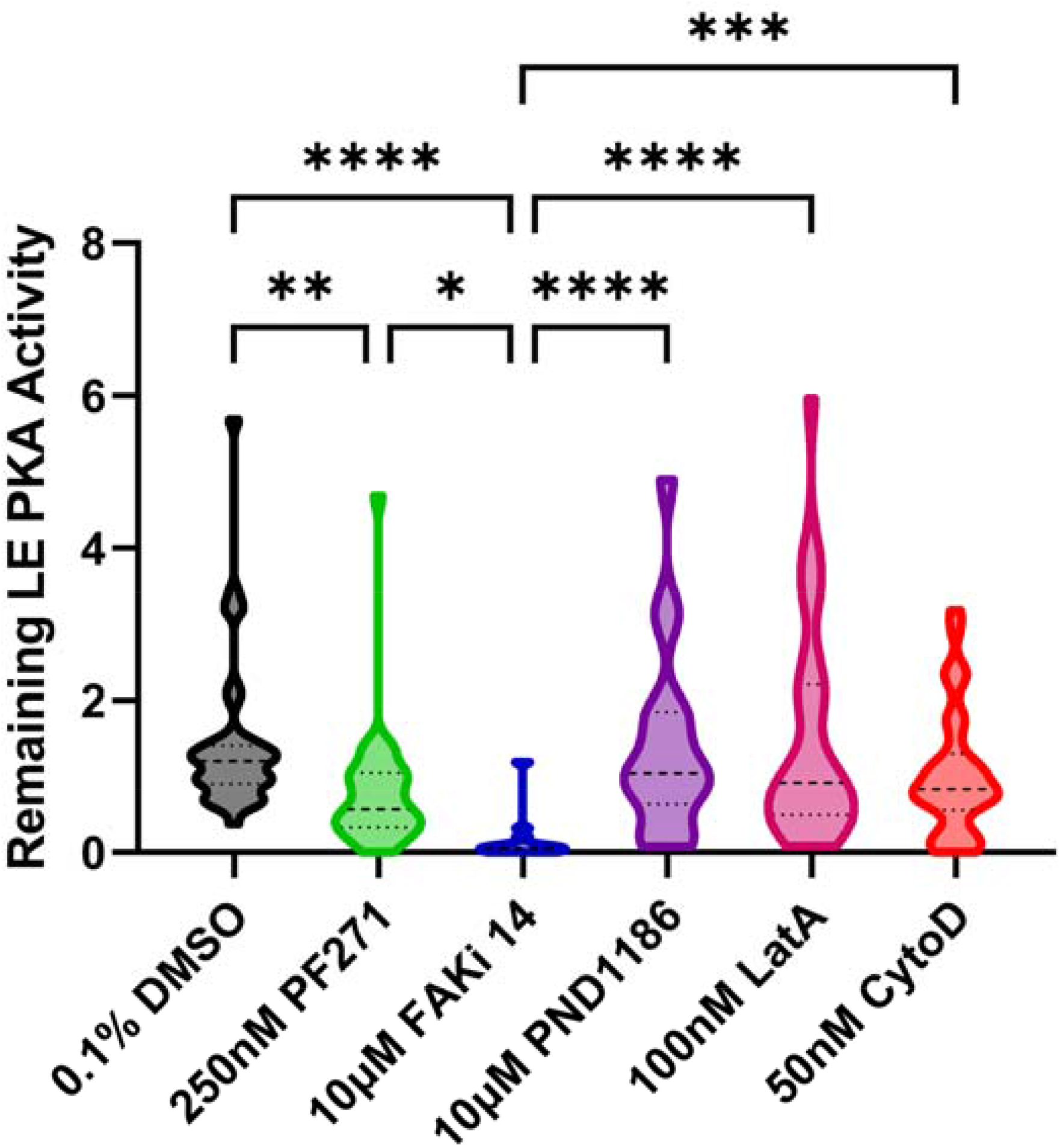
Comprehensive graph including all test results from Kruskal-Wallis assessment of the treatment of SKOV-3 cells with pharmacologic inhibitors. Comparisons shown in Figure 2 and Supplemental Figures 1-3 are a subset of all comparisons assessed by Kruskal-Wallis test, the complete results of which are shown here. * p≤0.05, ** p≤0.01, *** p≤0.001, **** p≤0.0001.

**Supplemental Figure 5:**
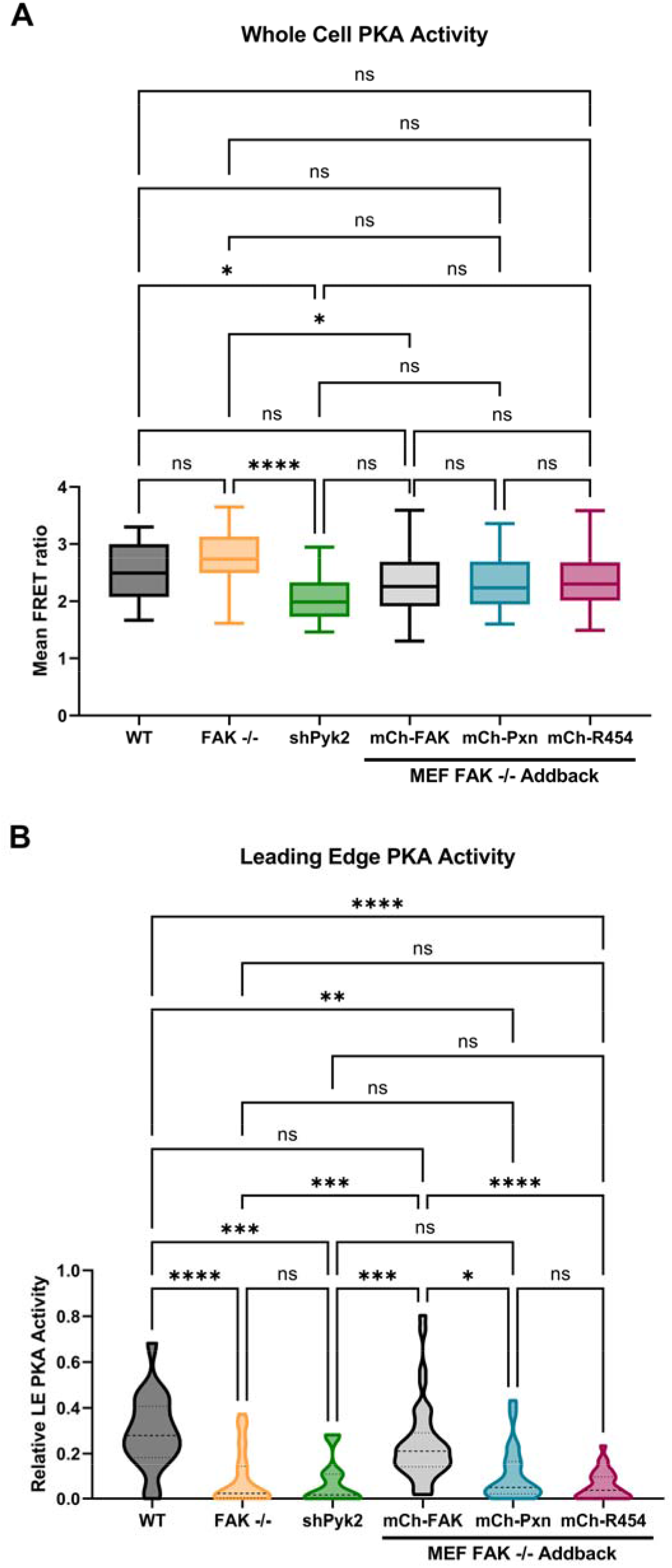
Comprehensive graph including all test results from Kruskal-Wallis assessment of MEF cell lines and MEF FAK-null cells expressing mCherry tagged addback constructs. Comparisons shown in Figures 3 and 4 are a subset of all comparisons assessed by Kruskal-Wallis test, the complete results of which are shown here. (A) Mean PKA activity as assessed by mean FRET ratio. (B) Relative leading edge PKA activity * p≤0.05, ** p≤0.01, *** p≤0.001, **** p≤0.0001.

## Acknowledgements

The authors thank John Patterson and Zoe Edmunds for technical assistance. This work was funded by National Institute of Health grants R01GM117490 and R01GM137611 (to AKH). Some of the live-cell microscopy was performed in the Microscopy Imagin Core supported by the University of Vermont Cancer Center, Lake Champlain Cancer Research Organization, and the UVM Larner College of Medicine.

